# *Candida albicans* infiltrates colon and rectal cancers causing therapeutic resistance and decreased survival

**DOI:** 10.1101/2025.09.19.677366

**Authors:** Dennis J. Grencewicz, Alexander Loncar, Sylvain Ferrandon, McKenzie Kreamer, Dipankor Chatterjee, Yogita Mehra, Aspen Carson, Rebecca Hoyd, Shiva Jahanbakhshi, Fouad Choueiry, Matthew Anderson, Martin Benej, Dustin E Bosch, Jiangjiang Zhu, Jinghai Wu, Aaditya Pallerla, Therese Bocklage, Martin McCarter, Ahmad Tarhini, Bodour Salhia, Christopher Moskaluk, Greg Riedlingeer, Song Yao, Ashiq Masood, Sheetal Hardikar, Mmadili Ilozumba, Cornelia M. Ulrich, Carlos H.F. Chan, Craig Shriver, Sagila George, Dinesh Pal Mudaranthakam, Michelle Churchman, Robert J. Rounbehler, Laura Chambers, David P. Carbone, Matthew Kalady, Nicholas Denko, Daniel Spakowicz

## Abstract

The microbiome is increasingly recognized as a modifier of cancer progression and therapy response, yet the role of intratumoral fungi remains poorly defined. Here, we identify *Candida albicans* colonization within human colorectal tumors as a predictor of reduced survival and impaired radiation response. Leveraging the Oncology Research Information Exchange Network (ORIEN) cohort, we show that high intratumoral *Candida* burden is associated with decreased survival across multiple gastrointestinal cancers, with the strongest treatment-specific effect in rectal cancer patients receiving radiotherapy. This observation was validated in independent rectal cancer cohorts using RNA sequencing and quantitative PCR.

In immune-competent murine colorectal cancer models, oral gavage of *C. albicans* resulted in intratumoral colonization, accelerated tumor growth, and radiation resistance, effects not observed with *Saccharomyces cerevisiae* or PBS controls. Colonized tumors exhibited increased hypoxia, altered metabolic and transcriptional programs, and distinct expression of genes linked to cytokine signaling and cell survival. Hypoxia conditioned *C. albicans* secreted metabolites that directly conferred radiation resistance to colorectal cancer cells *in vitro*, implicating a cancer cell intrinsic mechanism independent of immune signaling. Untargeted metabolomics revealed enrichment of nucleosides and lipid oxidation intermediates under hypoxia, suggesting that *C. albicans* metabolites may provide substrates facilitating tumor recovery after irradiation.

These findings establish *C. albicans* as a causal modifier of tumor biology and radiation response, highlighting intratumoral fungi as future potential therapeutic targets. Modulating fungal colonization or metabolism may improve radiotherapy outcomes and broaden our understanding of interactions between microbes and tumors.

## Introduction

The impact of the human microbiome on tumor development, growth and response to therapy has been a topic of significant recent research^1–3^. Effects driven by the microbiome can be detected at great distances from the gut including in the skin, breast, lung, brain and liver^1,4–10^. The concept that microbes travel to and inhabit anatomic sites distant from the gut and can produce small molecules that regulate host processes was first studied in the context of metabolic disorders^11^. This theory has since been introduced to oncology, now with robust evidence supporting that microbes in the gut and elsewhere can influence tumorigenesis and response to cancer treatments^1,3,12–20^. For instance, gut microbiota have been shown to modulate type I interferon expression in tumors through butyrate production, thus leading to impaired radiation effects^20^. Further, it is now apparent that microbes can become resident within tumors^4–6,21^, and that intratumoral microbes have the capacity to change the immune compartment of tumors^9^. Therefore, we propose that microbes that within tumors also have direct access to influencing tumor cells’ response to therapy. We hypothesize that cancerous tissues represent environments in which dynamic alterations of the microbiome can be readily established, particularly under conditions of metabolic dysregulation or radiation-induced epithelial injury.

Multiple studies have identified a relationship between specific microbes and treatment response in rectal cancer. Notably, in colon and rectal cancer (CRC), there is an increased abundance of the opportunistic pathogen fungus *Candida albicans*^5,6,22^. *Candida albicans* is a known commensal microbe in the mouth, gastrointestinal tract, genitourinary tract, and skin, and is commonly affiliated with oral, genitourinary, and systemic infections. In a recent large study, the presence of *C. albicans* DNA inside colorectal tumors independently predicted metastatic disease and decreased survival in patients with gastrointestinal tumors, and increased transcriptionally active *C. albicans* within in rectal tumors was associated with decreased survival^5^. Furthermore, previous work has separately shown that use of antifungal drugs can rescue radiation response in murine models of breast cancer orally supplemented with *C. albicans*^23^. An increase in Dectin-1 innate immune receptor signaling, a key sensor for fungi, was identified as the driver of decreased radiation response^23^. Although this Dectin-1 immune mechanism comments on potential systemic effects of *Candida* host colonization, it does not assess intratumoral tumor cell effects. Therefore, it remains unclear whether *C. albicans* directly modulates tumor cell biology upon intratumoral colonization, and how such colonization influences tumor progression and the response to cancer therapies, including irradiation.

Radiation resistance is a significant burden to treatment outcomes for cancer patients, including in the context of rectal cancer^24–31^. As a part of the standard of care for rectal cancer patients with advanced disease, radiation therapy is routinely administered^32^. However, a significant proportion of rectal cancer patients treated with chemoradiation will eventually have their tumors recur at local or distant sites, with potentially upwards of 35% of patients treated with radiotherapy and surgery developing distant metastases^33–36^. Prior studies have established that oxygen is integral to radiation-induced cell killing by facilitating DNA damage^37^. Therefore, in human tumors, pretreatment hypoxia has developed into an independent prognostic indicator of poor patient response to radiation therapy^37^. Hypoxia also contributes to tumor immune evasion by reducing immune cell infiltration and function^29–31,38,39^. In addition to making tumors radiation-resistant, hypoxia has also been shown to promote malignant progression in preclinical and clinical studies^26,39^.

The following study reports that intratumoral colonization by *Candida* species is associated with a decreased response to radiation and reduced survival in patients with human CRC. Further, using an immune-competent *in vivo* mouse model of CRC, we show that the oral gavage of *Candida albicans* causally confers radiation resistance and increased tumor growth. Additionally, tumors from mice inoculated with *C. albicans* exhibit altered immune infiltration, metabolism, and cell signaling. To attempt to understand the mechanism of enhanced tumor growth, we also studied the effects from *C. albicans* metabolites on cancer cells specifically, including untargeted metabolomics on *C. albicans* cultured in normoxic and hypoxic conditions to determine whether metabolites from *C. albicans* cultured in hypoxic conditions may contribute to a change in tumor growth. We observed that *Candida* exhibits cell-intrinsic and extrinsic effects on the tumor and its microenvironment, mediated by metabolites produced in hypoxic environments. The novel cell-intrinsic effect presented here offers deeper insight into how multifaceted host-*Candida* interactions drive worsened cancer outcomes. These findings support that the presence of *C. albicans* within human tumors can causally modify tumors, which may play a larger role in decreased therapy response and colorectal cancer patient survival than previously appreciated.

## Methods

### Human Study Design

The Oncology Research Information Exchange Network (ORIEN) is an alliance of 18 US cancer centers established in 2014. All ORIEN alliance members utilize a standard Total Cancer Care® (TCC) protocol. As part of the TCC study, participants agree to have their clinical data followed over time, to undergo germline and tumor sequencing, and to be contacted in the future by their provider if an appropriate clinical trial or other study becomes available. TCC is a prospective cohort study with a subset of patients enrolled in the ORIEN Avatar program, which includes research-use-only (RUO)-grade whole-exome tumor sequencing, RNA-sequencing, germline available under aCC-BY 4.0 International license. M2GEN, the commercial and operational partner of ORIEN, harmonizes all abstracted clinical data elements and molecular sequencing files into a standardized, structured format, enabling the aggregation of de-identified data for sharing across the network.

### Human RNA-Sequencing Methods

ORIEN Avatar specimens undergo nucleic acid extraction and sequencing at HudsonAlpha (Huntsville, AL) or Fulgent Genetics (Temple City, CA). For frozen and optimal cutting temperature (OCT) tissue RNA extraction, the Qiagen RNAeasy plus mini kit is used, generating 216 bp average insert size. For formalin-fixed paraffin-embedded (FFPE) tissue, Covaris Ultrasonication FFPE DNA/RNA kit is utilized to extract both DNA and RNA, generating a 165 bp average insert size. RNA-sequencing (RNA-seq) is performed using the Illumina TruSeq RNA Exome with single library hybridization, cDNA synthesis, library preparation, and sequencing (100 bp paired reads at Hudson Alpha, 150 bp paired reads at Fulgent) to a coverage of 100M total reads/50M paired reads. RNA-seq tumor pipeline analysis is processed according to the workflow outlined below, using the GRCh38/hg38 human genome reference sequence and GenCode build version 32. Adapter sequences are trimmed from the raw tumor sequencing FASTQ file. Adapter trimming via k-mer matching is performed in conjunction with quality trimming and filtering, contaminant filtering, sequence masking, guanine-cytosine (GC) content filtering, and entropy filtering. The trimmed FASTQ file is used as input to the read alignment process. The tumor adapter trimmed FASTQ file is aligned to the human genome reference (GRCh38/hg38) and the GenCode genome annotation v32 using the STAR aligner. RNA expression values are calculated and reported using estimated mapped reads, fragments per kilobase of transcript per million mapped reads (FPKM), and transcripts per million mapped reads (TPM) at both the transcript and gene levels based on transcriptome alignment generated by STAR. For model murine tumors, we used Powerfecal pro kits from Qiagen to extract RNA and DNA. RNA-seq analyses utilized Deseq2 to determine differential gene expression, FGSEA to highlight pathway analysis, and {tmesig} for determining gene signatures from Deseq2 results.

### Microbial Culture

Several strains of fungi were cultured for murine inoculation and *in vitro* experiments, including *Candida albicans* SC5413 (ATCC MYA-2876), and *Saccharomyces cerevisiae* 1187. Overnight cultures for each strain were struck out from −80 °C freezer stocks on yeast peptone dextrose (YPD) solid agar. Individual colonies were picked and cultured in liquid broth at 37 °C for 16-20 hours using liquid YPD media to achieve log phase growth, measured by generating fungal growth curves. Cultures were shaken at 120rpm after singular colony inoculation from plated fungi into 45 mL of media in a 50 mL conical centrifuge tube. Overnight cultures were counted using a hemocytometer. For murine gavage, 1×10^7 microbes were gavaged in a total of 200 uL of PBS.

To generate normoxic and hypoxic conditioned media, liquid culture of *C. albicans* was grown in 37 °C room air conditions or within an anaerobic chamber with oxygen less than 120 ppm (Coy Chambers) over 16-20 hours. After overnight culture, cultures were spun at 1200 G for 12 minutes to create a cell pellet. Supernatant was eluted into a fresh 50 mL Falcon tube and then syringed through a 0. 22 µM filter to create a sterile supernatant. Sterile conditioned media was made fresh and stored for less than one week at 4 °C in room air before use.

### Murine Models

Animals were purchased from Charles River (housed in groups of 5) and were injected subcutaneously on the right flank with 0.5×10^6 MC38 cells with tumor growth measured by calipers using the formula (L×W2)/2. When tumors reached approximately 250 mm^3^, we randomized mice into three groups and gavaged the mice with 200 uL phosphate buffered saline alone (PBS, control) or with 1×10^7 cells of either *Candida albicans* (CA, test) or *Saccharomyces cerevisiae* (SC, control) for the one-time gavage experiment. For the multiple-gavage experiment using MC38 cells, cells were implanted, and four gavages took place every other day, starting 4 days after cell implantation. MC38 tumors were irradiated 48 hours after the last gavage with image guidance on a small animal radiation research platform (SARRP) using a dose of 7.5 Gy. The experiment was ended six days later when tumor sizes reached early removal criteria. Untreated tumors were typically harvested after sacrifice, when most tumors were at the removal criteria. Mice’s stool was collected before tumor harvesting, and the tumor, tumor-draining lymph node, liver, spleen, colon, and blood were harvested upon sacrificing the mice.

### *In Vitro* Experiments

*Candida albicans* (strain SC5314) was in YPD until cells reached log phase growth (16-20 hours at 37 °C) in either normoxic or hypoxic conditions as described above. *In vitro* radiation response was assessed by plating 2×10^3 MC38 into 6-well plates with 33% conditioned media. The next day, cells are irradiated with 2 to 10 Gy for the clonogenic survival assay and incubated for eight days. Colonies are counted using a crystal violet assay, and plating efficiency was calculated based on comparing the 0 Gy colony number to increasing doses of radiation.

### Metabolomics

We have used the Thermo LC-Q-Exactive Hybrid MS system on samples of *C. albicans* cultures grown *in vitro* in either 21 or 1% oxygen in rich YPD media for 16 hours until the culture was in log growth phase. Cell pellets and media were collected and normalized by cell number for LC-MS/MS quantification for water-soluble and insoluble metabolites and lipids. Samples were extracted with either methanol or isopropanol, dried, resuspended in either 50/50 water-methanol or acetonitrile, and spiked with stable isotope-labeled internal standards for quality control.

### Quantitative PCR

Nucleic acids were extracted from FFPE tissue samples of the Iowa cohort as previously described (QIAGEN FFPE DNA Advanced UNG)^40^. qPCR amplification with SYBR green (Applied Biosciences master mix) was performed on a QuantStudio 3 instrument in triplicate with three pairs of *Candida* genus-specific primers. Melt curves were assessed with comparison to amplicons from a control *C. albicans* strain ATCC 18804. Detection was considered amplification within 30 cycles and a characteristic amplicon melt curve for at least 2 of the 3 primer pairs.

## Results

### *Candida* can be identified within colorectal human tumors and increased abundance is associated with decreased survival

Given that fungi in the genus *Candida* have previously been linked to changes in survival in several cancer types^5,41–43^, we wanted to determine whether this relationship may also be related to response to therapy and whether we could identify a stronger causal driver of this relationship. Leveraging a patient cohort of over 17,000 tumors from Oncology Research Information Exchange Network (ORIEN) cohort, which has access to robust treatment, outcome, and transcriptomics data for each patient, we determined the intratumoral microbiome of this cohort using {exotic}, a tool capable of carefully identifying microbial sequences from bulk RNA-sequencing datasets (Fig. 1A)^44^. We first asked whether intratumoral *Candida* was associated with overall survival in several cancer types where *Candida* has been previously associated, including head and neck squamous cell carcinoma (Fig. 1B), esophageal (Fig. 1C), gastric (Fig. 1D), pancreatic (Fig. 1E), hepatic (Fig. 1F), other GI tumors (Fig. 1G), and colorectal cancer (CRC) (Fig. 1H). Given sequencing depth, we identified microbes most confidently at the genus level. When stratifying patient samples by “high” versus “low” *Candida*, we observed that patients with high *Candida* had a significantly decreased overall survival.

**Figure 1.**
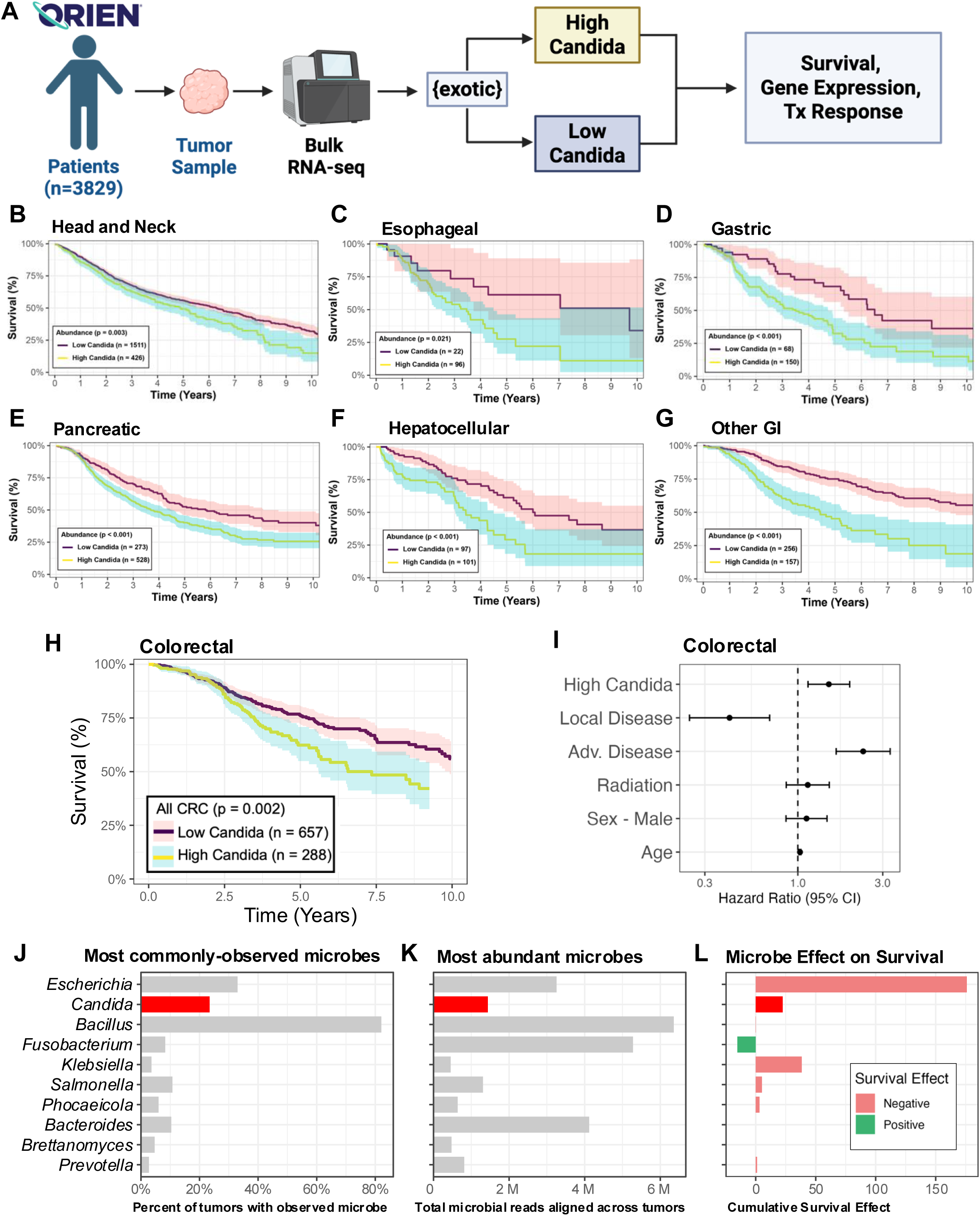
Intratumoral *Candida* associates with survival across several cancer types. **A)** Summary of the ORIEN cohort and the method for evaluating microbial reads from tumor RNAseq. **B-H)** The percent of patients alive over time when stratified by tumors containing high *Candida* across several cancer types. **B)** Head and Neck, **C)** Esophageal, **D)** Gastric, **E)** Pancreatic, **F)** Hepatocellular, **G)** Other GI, **H)** Colorectal. **I)** *Candida* significantly associates with overall survival in CRC when controlling for confounding variables. **J)** Microbes observed in ORIEN RNAseq data after {exotic} pipeline alignment, ordered according to the fraction of tumors in which reads aligned to that microbial genus. **K)** The most abundant microbes observed in ORIEN CRC samples, ordered by the total number of reads. **L)** Cumulative survival effect from particular microbes on ORIEN CRC patients.

Because *Candida* is a known opportunistic pathogen that can colonize the GI tract, we focused our following analyses on CRC (Fig. 1H) due to the increased likelihood of subsequent tumoral colonization, previous literature precedent^5,41^, as well as the depth of patient data available for the ORIEN cohort samples. Sub-setting to ORIEN patients with only primary CRC (Table 1), the effect on survival observed in samples with high intratumoral *Candida* burden remained after controlling for clinical stage, age, sex, and treatment (Fig. 1I). We then used {exotic} to more broadly qualify microbes within these CRC samples based on prevalence (Fig. 1J), rank order abundance (Fig. 1K), and overall impact on survival^45^ (Fig. 1L). *Candida* was one of several microbes with increased abundance and prevalence in tumors from the cohort, with other reported intratumoral microbes including those in the genera *Escherichia*, *Bacillus*, and *Fusobacterium* also observed (Fig. 1J-L). These data supported that *Candida* could reliably be identified in the ORIEN cohort CRC samples and its increased prevalence was associated with decreased survival, corroborating previous data^5,6,41^.

Next, to parse out whether the change in overall survival associated with increased intratumoral *Candida* could be secondary to changes in therapeutic response, we subset the ORIEN cohort to patients with primary rectal cancer that received radiation treatment (Fig. 2A). In this cohort, high *Candida* burden again predicted decreased survival (Fig. 2A) with disease stage, sex, BMI, and age controlled for (Fig. 2B). To confirm our ability to identify *Candida* within tumors and understand whether intratumoral *Candida* burden was affected by radiation itself, we developed two validation cohorts of human colorectal and rectal cancer samples. We first completed bulk RNA-sequencing on matched patient pre-irradiated biopsies and post-irradiation rectal tumor resections from an Ohio State University cohort (Table 2, Fig. 2C). Intratumoral *Candida* burden was assessed with {exotic} to assess whether irradiation itself affected the capability of *Candida* to invade tumors. We saw no significant difference in *C. albicans* RNA-seq counts based on whether samples were taken before or after irradiation (Fig. 2D), suggesting that *Candida* can colonize and inhabit a tumor at a similar measurable density irrespective of whether the tumor previously received local radiotherapy.

**Figure 2.**
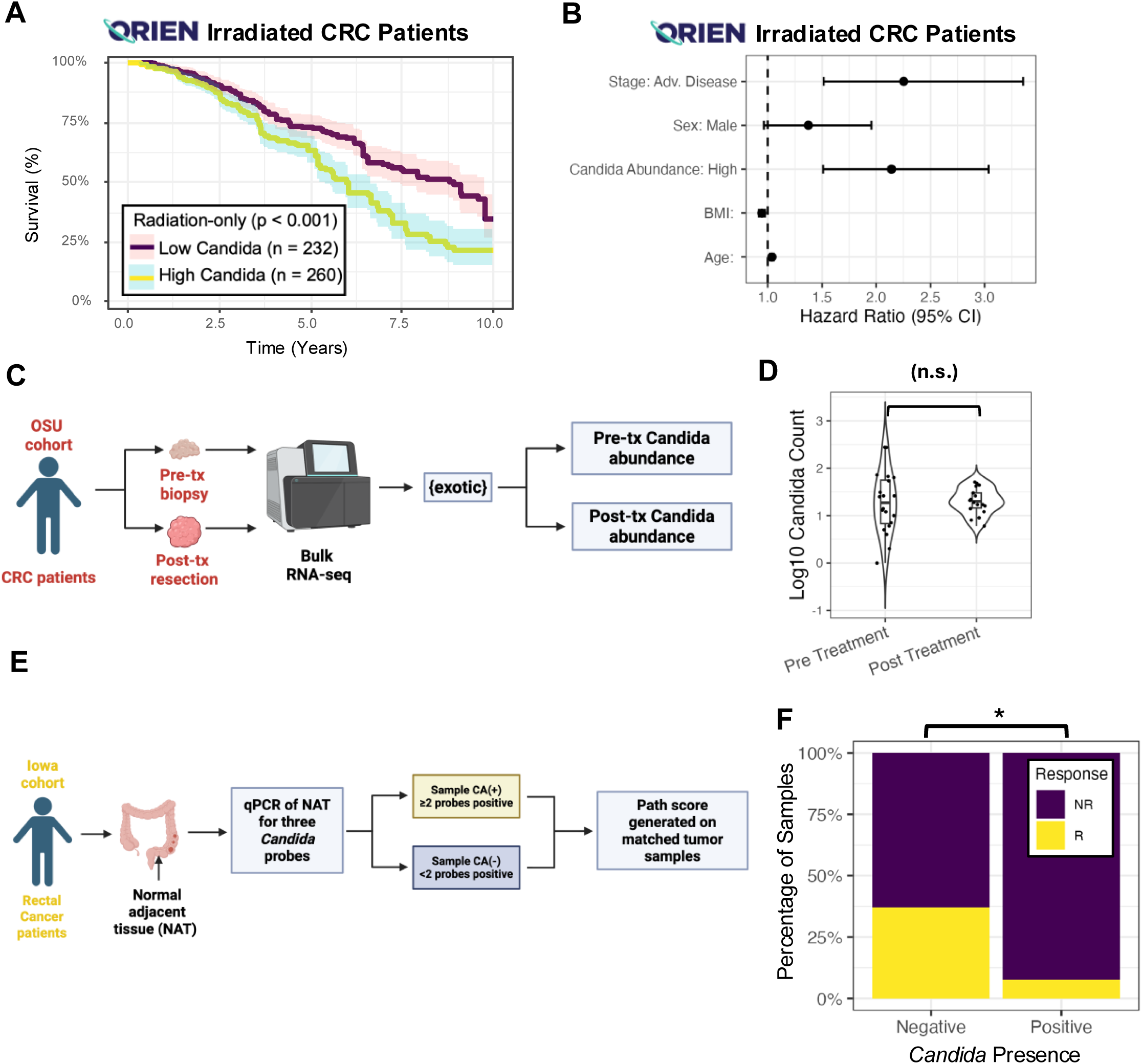
*Candida* tumor colonization is unaffected by radiation and *C. albicans* presence in tumor adjacent normal tissue can predict response. **A)** *Candida* most strongly associates with overall survival in the subset of ORIEN CRC patients who received radiation treatment. **B)** The association of *Candida* with overall survival remains significant in the CRC cohort of patients treated with radiation even when controlling for confounding factors.**C)** Summary of the OSU validation cohort and the method for evaluating microbial reads from tumor RNAseq. **D)** RNAseq counts of aligned to *Candida* in OSU cohort, with no significant difference observed between samples taken before (n=20) or after (n=18) radiation therapy.**E)** Summary of the Iowa validation cohort and the method for identifying Candida in tumor-adjacent normal (TAN) samples. **F)** Percentage of *Candida* negative (n=30) and positive (n=10) rectal normal adjacent tissue samples in the Iowa cohort in which patients showed pathological response to radiation. * p < 0.05.

To determine whether *Candida* could be found in human tissue samples beyond using RNA-sequencing-based techniques, we used three quantitative PCR (qPCR) probes specific for the *Candida* genus to assess *Candida* burden in rectal cancer tumor-adjacent normal (TAN) samples in a third cohort from University of Iowa (Table 3, Fig. 2E)^40^. Tumors were deemed “*Candida* positive” if at least two genus-specific amplicons were detected. Further, using clinical information on pathological treatment response scores at time of resection, we identified that *Candida*-positive status in rectal TAN samples could predict response to radiation (p<0.05) (Fig. 2F). Together with the association of high intratumoral *Candida* and decreased survival, this finding suggests that mucosal colonization by *Candida* may reduce tumor response to radiation, decreasing overall survival.

### Rectal tumors with increased *Candida* exhibit changes in gene expression

After observing the effects on survival and therapy response from intratumoral *Candida*, we used the bulk RNA-sequencing available with ORIEN samples for hypothesis generation about how intratumoral *Candida* may be contributing to this phenotype. We completed differential gene expression and pathway analysis on samples deemed high or low *Candida* in the ORIEN rectal cancer patient cohort that received radiation (Fig. 3A). We observed a significant increase in noncoding RNA expression in the high-*Candida* tumors, specifically RNAs in the RN7SL and RNU5 families. Further, changes in SMAD and SNORA expression were noted in high-*Candida* tumors (Fig. 3A). Several genes were also upregulated in the low-*Candida* tumors, including genes peripherally associated with the Notch signaling pathway (Fig. 3A). Subsequently, we completed FGSEA pathway analysis and found that in rectal tumors with high *Candida,* oxidative phosphorylation, targets of MYC, targets of E2F, MTORC1 signaling, protein secretion, and DNA repair pathways were enriched (Fig. 3B). Further, WNT/b-Catenin, apical junction, coagulation, inflammatory response, pancreatic beta cell, TNFa signaling by NFKb, and KRAS signaling pathways were enriched in patients with low intratumoral *Candida* (Fig. 3B). Using immune deconvolution to better understand the potential interactions of the immune system as a phenotypic driver of radiation resistance in high *Candida* tumors, we did not see robust changes in the immune compartment, including no significant changes to innate immune signaling and particularly no changes to Dectin-1 signaling targets or intermediates (Fig. 3C). Transcriptional analyses of these tumors suggested that there may be an effect of *Candida* on cancer cell transcriptional regulation by noncoding RNAs, the innate and adaptive immune systems, as well as metabolic programming, beyond the reported effect on survival from increased Dectin-1 signaling in hosts colonized with *Candida*^23^.

**Figure 3.**
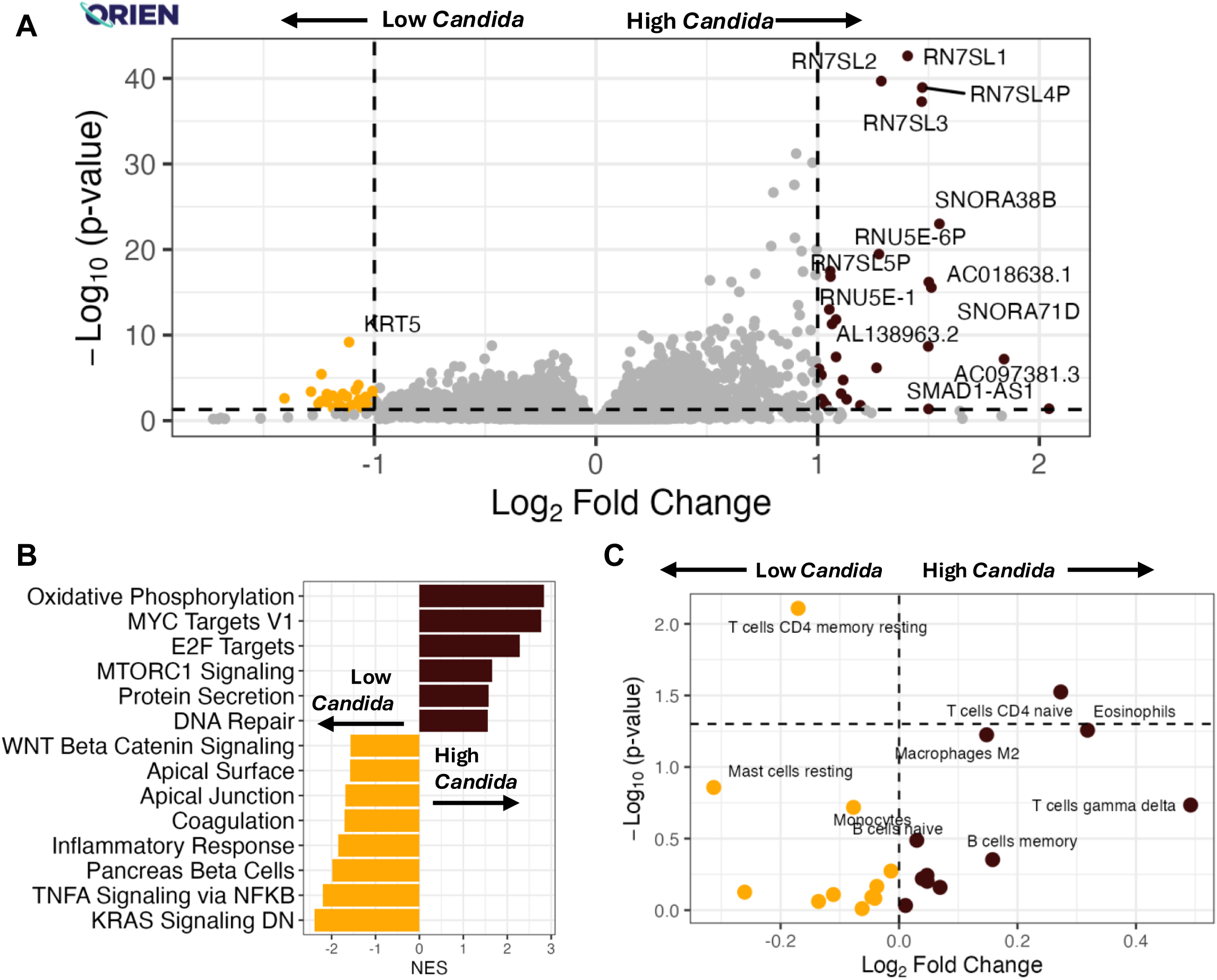
Patient CRC tumors with high *Candida* have altered gene expression. **A)** Differential gene expression between ORIEN CRC tumors with low versus high *Candida*. **B)** FGSEA Pathway analysis from the Molecular Signatures Database (MSigDB) on ORIEN human CRC samples from the ORIEN CRC Cohort (Fig. 1) showing tumors with high *Candida* have increased oxidative phosphorylation and cell signaling pathways and decreased immune activation pathways.**C)** Immune deconvolution analysis from ORIEN human CRC samples showing that tumors with high or low *Candida* differ in gene expression related to naïve and memory CD4 T Cells but no other immune cell types.

### Orally-gavaged *Candida albicans* colonizes tumors and affects growth rate and radiation response, gene expression, and intratumoral hypoxia in murine models

After observing a strong relationship between intratumoral and peritumoral *Candida* with decreased survival in CRC patients as well as changes to the transcriptional regulation of rectal tumors colonized with higher levels of *Candida*, we looked to assess whether CRC colonization with *Candida* directly caused these phenotypes using a mouse model of CRC. Syngeneic MC38 murine CRC cells were inoculated into the flank of male immunocompetent C57BL/6 mice. After tumor implantation, mice were orally gavaged with either *C. albicans*, *Saccharomyces cerevisiae* (SC, fungal gavage control), or PBS. Two days after gavage, mice were then treated with one dose of 7.5 Gy radiation, and tumor volumes were tracked until experimental endpoint (Fig. 4A).

**Figure 4.**
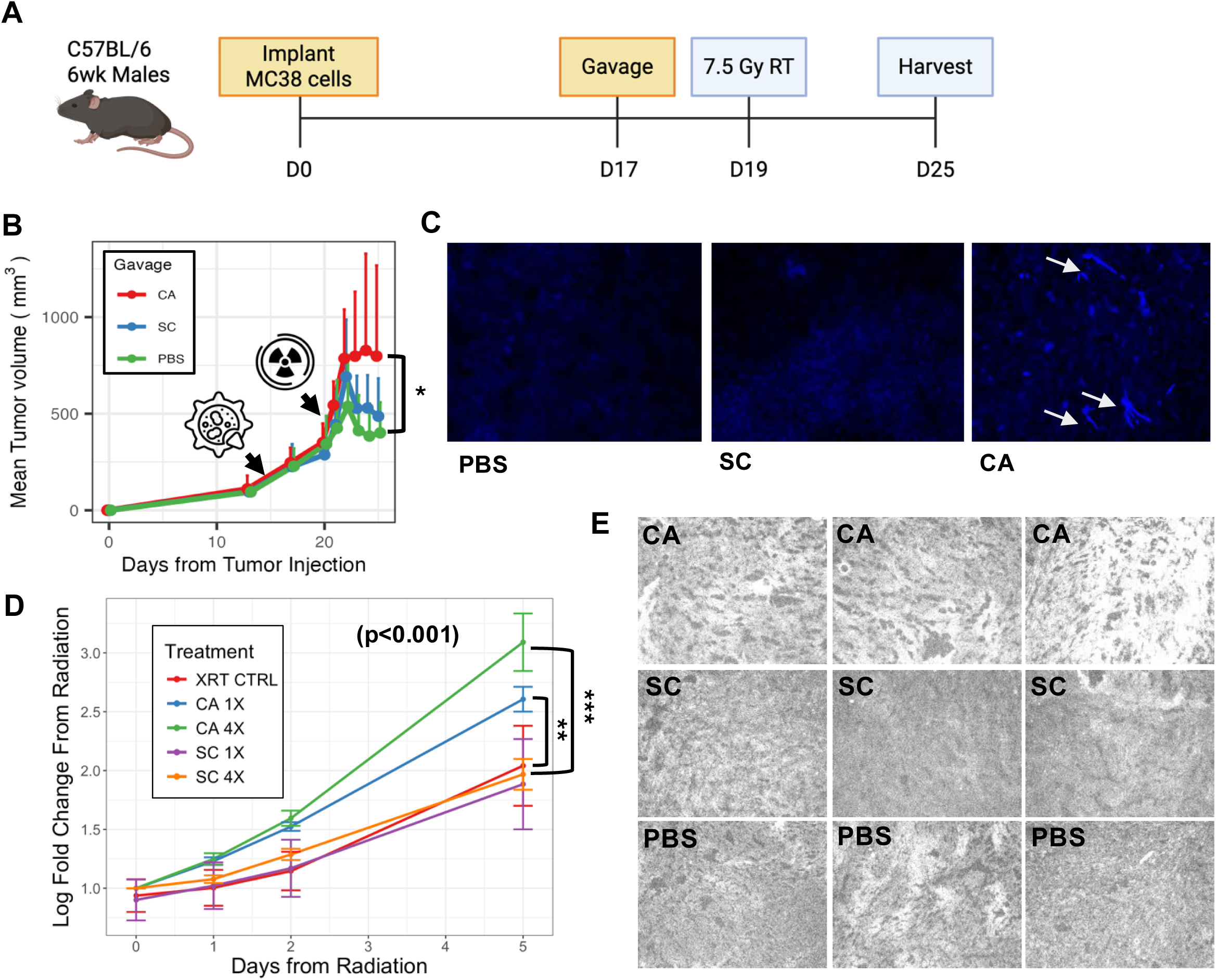
*Candida albicans* colonization of CRC mouse tumors causes radiation resistance. **A)** Schema of heterotopic MC38 CRC murine model treated with oral gavage of fungi before radiation therapy. **B)** Oral gavage of CA causes decreased response to radiation in murine CRC tumors compared to SC and PBS gavage. **C)** Fungal staining with calcofluor white shows intratumoral hyphae in tumors from mice that were gavaged with CA with no positive staining in SC or PBS conditions. **D)** Increasing dose of oral gavage of CA causes decreased response to radiation in murine CRC tumors compared to SC and PBS gavage. All mice received one dose 7.5 Gy irradiation. XRT CTRL (n=6), mice received PBS gavage with no addition of fungi; CA 1X (n=6), one gavage CA; CA 4X (n=6), four gavages, SC 1X (n=6), one gavage SC; SC 4X (n=6), four gavages SC **E)** Gavage of CA 4X leads to increased intratumoral hypoxia per pimonidazole immunofluorescence staining from murine tumors after CA 4X, SC 4X, or PBS 4X gavage and radiation. All images taken at 40x and pimonidazole inverted to greyscale with increasing intensity represented as white. ** p < 0.01, *** p < 0.001.

Mice gavaged with *C. albicans* (CA) showed significantly increased tumor volume compared to SC-gavaged mice and the PBS-gavaged group (Figure 4B). The mean tumor volume of SC-gavaged mice was smaller than the CA-gavaged group. (Figure 4B). Upon necropsy, tumors were sterilely removed, flash frozen, sectioned, stained with the pan-fungal stain calcofluor white (CFW), and imaged. Fungal hyphae were visible in the tumors of CA-gavaged mice but not SC or PBS-gavaged mice (Figure 4C). Given that these mice were SPF and fungal-depleted, this demonstrated that the orally-gavaged CA translocated to the heterotopic flank CRC tumors.

We conducted a second heterotopic syngeneic mouse experiment using the MC38 CRC model to assess whether CA gavage affected radiation response in a dose dependent manner by comparing one-time gavage (1X) to 4-times gavages (4X) (Fig, 5B). When comparing 1X to 4X gavage of CA, mice gavaged four times had significantly increased tumor volume after irradiation compared to a one-time gavage (Fig. 4D). Further, one-time gavaged mice still maintained increased tumor volumes after irradiation compared to either four- or one-time SC gavaged mice or PBS gavaged mice (Fig. 4D). Before euthanasia, mice were injected with pimonidazole, a 2-nitroimidazole compound used as a hypoxia marker to identify poorly oxygenated regions in tissues. We observed gavage-dependent increases in pimonidazole staining with the CA four-time gavage group exhibiting the strongest pimonidazole staining (Fig. 4E), suggesting this condition was the most hypoxic.

Given the effect on tumor volume after radiation therapy, tumors were submitted for bulk RNA-sequencing to the Applied Microbiology Services Lab (AMSL) at The Ohio State University, to assess their transcriptomic profile to determine the cause of radioresistance in this model system. Differential gene expression analysis was completed with Deseq2 to determine changes in gene expression between tumors harvested from mice gavaged with CA, SC, or PBS. Comparing tumors from mice gavaged with CA versus PBS, CA-gavaged murine tumors exhibited upregulation of cytokine signaling genes including Osm, Il3ra2, and Ucn2 (Fig. 5A). Comparing tumors from SC-gavaged mice and CA-gavaged mice, increases in Abcd2, Asb2, and Dapk1 were observed in CA-gavaged mice (Fig. 5B). After determining the differentially expressed genes in these mouse tumors, we looked for changes in gene signature pathways from the MSigDB Hallmark database. In the PBS-gavage condition relative to CA condition, the adipogenesis, oxidative phosphorylation, DNA repair, inflammatory response, unfolded protein response, and IL6 signaling were all increased in abundance (Figure 5C). Comparing the SC-gavage condition to the CA condition, the myogenesis, adipogenesis, oxidative phosphorylation, fatty acid metabolism, and TNFa signaling pathways were all overrepresented (Figure 5D).

**Figure 5.**
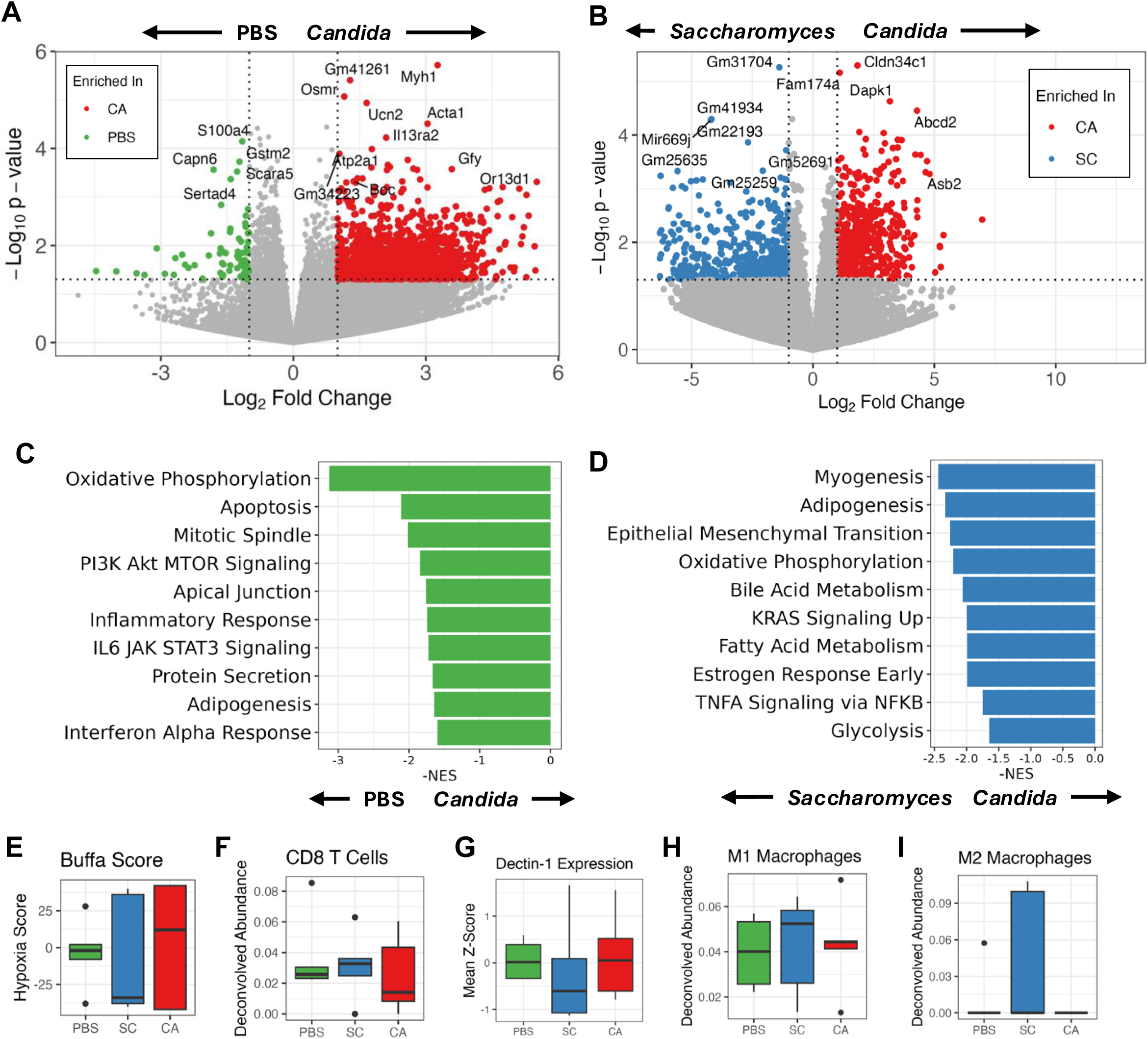
Transcriptional programming of murine tumors affected by intratumoral*Candida albicans*. **A-B)** Volcano plot representing differentially expressed genes between tumors from mice gavaged between **A)** CA and PBS, and **B)** CA And SC analyzed in DeSeq2. **C-D)** FGSEA pathway analysis comparing enriched pathways from CA gavaged murine tumors compared to **C)** PBS or **D)** SC. **E-I)** Targeted RNAseq gene signature analysis using tmesig for **E)** Buffa hypoxia scores, **F)** CD8 T Cells, **G)** Dectin-1 signaling, **H)** M1 macrophages, and **I)** M2 macrophages.

Based on the genes and pathways we observed to be differentially regulated in CA-gavaged mice, we became interested in testing for differences in specific gene signatures related to immune activation and tumoral hypoxia. We found that Buffa hypoxia^46^ score was elevated (Fig. 5E) and that CD8 T cells were decreased (Fig. 5F) in the CA condition compared to the SC and PBS conditions, but these findings did not reach statistical significance (p>0.05). Further, we did not observe differences between Dectin-1 signaling (Fig. 5G), M1 macrophages (Fig. 5H), and M2 (Fig. 5I), suggesting no overall changes in innate immune activation within the murine tumors.

These murine model data support that oral gavage of *C. albicans* in syngeneic, SPF mouse models of CRC lead to intratumoral colonization of CA, modulation of transcriptional programs related to metabolism, increased tumoral hypoxia, and decreased response to radiation therapy. Due to these data suggesting a novel a cancer cell-intrinsic effect of *C. albicans,* potentially modified by hypoxia, we next considered whether molecules or metabolites secreted by *C. albicans* facilitate this radioresistant phenotype.

### *Candida albicans* metabolites directly modify cancer cell survival and radiation resistance

To study the effect of secreted molecules from *C. albicans* on cancer cells, we cultured *C. albicans* in normoxic (21% oxygen) and hypoxic (0% oxygen in Coy Chamber) conditions to generate conditioned media. After centrifugation and filtration to remove *Candida*, the sterile conditioned media (CM) was applied to MC38 CRC cell lines for 12hrs. Then, MC38 cells underwent a clonogenic survival assay using 0, 2, 4, 6, 8, or 10 Gy irradiation (Fig. 6A). We observed significant increase in colony number after irradiation in the hypoxic CM condition, specifically in the 6, 8, and 10 Gy conditions (Fig. 6B). A clonogenic survival assay also showed that MC38 cells cocultured with hypoxic CM exhibited significant radiation resistance compared to normoxic CM and control media (YPD) as radiation dose increased (p < 0.05) (Fig. 6C).

**Figure 6.**
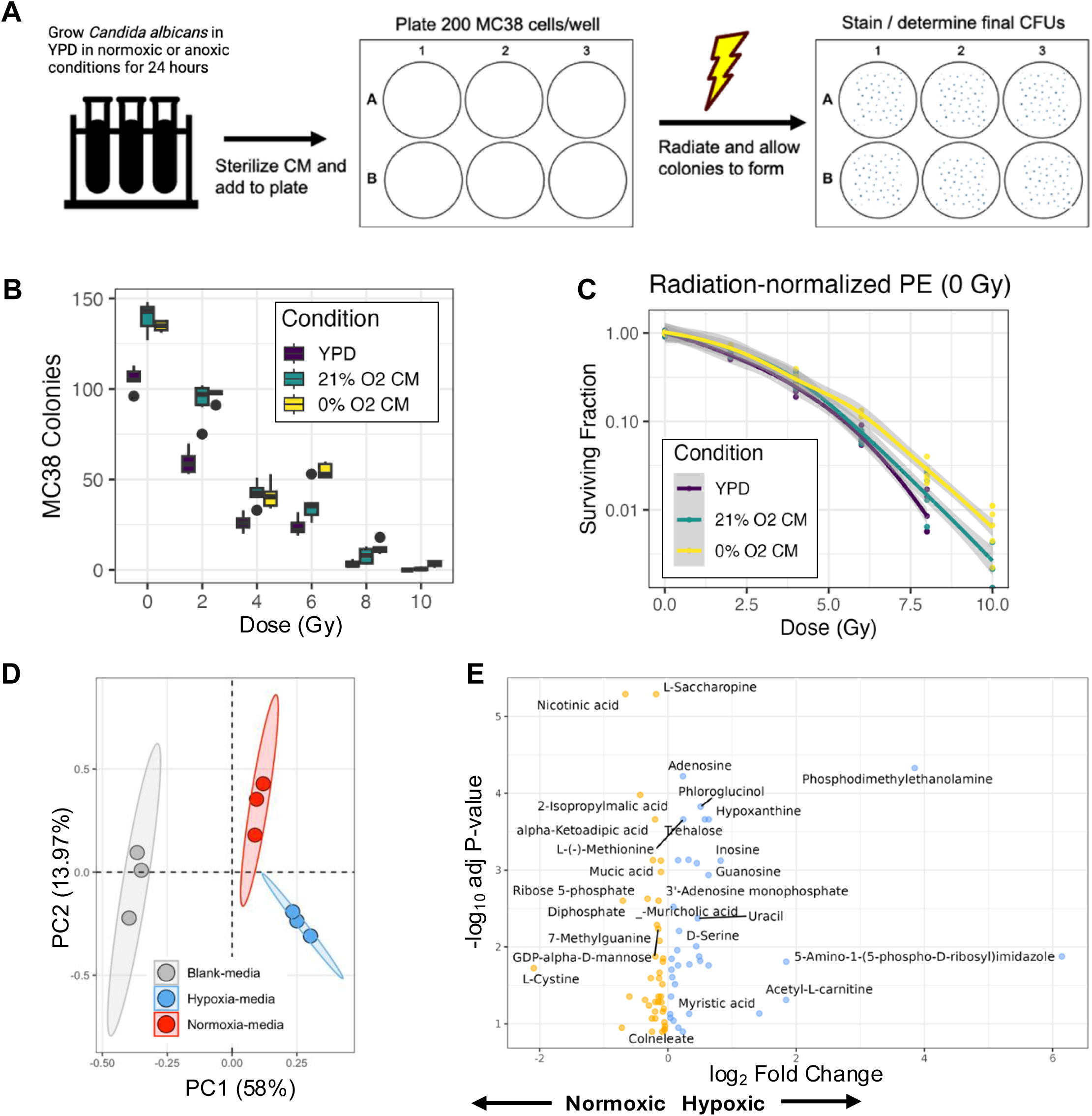
*Candida albicans* secretes molecules that confer radiation resistance in cell lines. **A)** Schema for culture of *C. albicans* in normoxic and anoxic conditions to generate sterile conditioned media (CM) for use in radiation response experiments with MC38 cells. **B)** MC38 colony number after clonogenic survival assay comparing anoxic CA CM to normoxic CM and media controls. **C)** Clonogenic survival assay shows increasing MC38 radiation resistance with addition of CA anoxic CM compared to normoxic CM and control YPD media. **D)** Principal Component Analysis of normoxic and hypoxic CA media after untargeted metabolomics shows unique metabolites in hypoxic and normoxic media. **E)** Differentially abundant metabolites in hypoxic and normoxic CA CM using untargeted metabolomics via LC-MS/MS quantification.

To identify whether *C. albicans* impacts tumor growth through the secretion of a metabolite, we cultured CA in normoxic and hypoxic conditions and measured intracellular and secreted metabolites via untargeted metabolomics. Principle component analysis significantly differed between high and low oxygen conditions (Fig. 6D). While we observed increases in glycolytic intermediates in the hypoxic CM similar to what we find reported for higher eukaryotes (such as elevated glucose 6P), we also found a decrease in the levels of both saturated and non-saturated fatty acids. Interestingly, the reduction fatty acid of hypoxic condition is accompanied by an increase in acetyl- and propionyl-carnitines (Figure 6E). In addition to the interesting findings regarding changes in lipid metabolism in *C. albicans* cultured in hypoxic conditions, there are notable differences in purine synthesis and excretion as well (Figure 6E). Adenosine, guanine, guanosine, inosine, and uracil are all seen with more relative frequency in the media of *C. albicans* cultured in hypoxic conditions (Figure 6E). Further, differences in fatty acid products are observed as well, with increased levels of carnitines secreted in hypoxic CA culture (Figure 6E).

## Discussion

Radiation resistance remains a major barrier to effective cancer care in many cancer types, including rectal cancer. A known limitation to the efficacy of radiotherapy is an increase in tumor hypoxia, which limits the formation of reactive oxygen species formation and subsequent DNA damage within tumors. This study identifies that increased intratumoral *Candida albicans* burden is associated with decreased radiation response and decreased overall survival in patients with rectal cancer. Further, we show that *C. albicans* can cause radiation resistance in mouse models of CRC, and that there is a cancer cell intrinsic effect of *C. albicans* conditioned media that facilitates radiation resistance.

Data from our mouse model show that oral gavage of *C. albicans* in mice bearing heterotopic colorectal tumors leads to decreased radiation response after local radiotherapy when compared to mice gavaged with *Saccharomyces cerevisiae* or PBS. Through immunofluorescent staining of tumor tissue with the calcofluor white stain, which binds to fungal cell wall components, we see that *C. albicans* has the capacity to migrate from the gastrointestinal tract to flank heterotopic tumors. These data also support that *C. albicans* can survive radiation therapy and replicate within tumors for at least 9 days after initial oral gavage. Further, by staining these tumors with pimonidazole to assess intratumoral hypoxia, murine tumors colonized with *C. albicans* showed increased levels of hypoxia, suggesting a potential relationship between the presence of *C. albicans* and hypoxic tumor development. Our bulk RNA-sequencing data did not exhibit changes in Dectin-1 signaling, which was previously suggested as the primary mechanism in models of breast cancer radiation resistance influenced by *C. albicans*^23^. This may have been related to when the tumors were harvested as opposed to a lack of Dectin-1 effect in the model, which remains undetermined but a direction of future study.

Overall, our mouse model supports a causal relationship between the presence of *C. albicans* within a tumor and a decreased response to radiotherapy. Noteworthy details of this model include that MC38 cells have a high microsatellite instability phenotype (MSI-High), indicating that this cell line has an increased tumor mutational burden. However, only ∼15% of rectal cancer are MSI-High; therefore, these data may not be representative of all genomic variants of rectal cancer. Future experiments plan to leverage several other mouse models, including several other cell lines and an orthotopic model of rectal cancer in immunocompetent mice. We also plan to reassess the potential dual phenotype of cell intrinsic and extrinsic effects of *C. albicans* by determining differences in immune signaling and cellular responses at several time points relative to radiation treatment.

To determine if the phenotype of increased tumor volume after radiation therapy in mice orally gavaged with *C. albicans* was mediated by secreted or expelled metabolites, we completed untargeted metabolomics on *C. albicans in vitro* culture that occurred in normoxic and hypoxic conditions. Given the heterogeneity within the microenvironment of tumors and the association with increased hypoxia in tumors harboring *C. albicans*, we wanted to further understand if *C. albicans* in hypoxic environments may be producing metabolites that could further tumor progression and treatment resistance in a positive feedback loop leading to worsened irradiation response. *Candida albicans* is well studied to phenotypically switch between hyphal and yeast forms based on its environment^48–50^, with one established hyphal driver of *C. albicans* being hypoxia^51^. Previous data from our group also supports that microbes can adapt their transcriptional profile based on their within tumors^47^. We identified that *C. albicans* grown in hypoxic environments produces a unique set of metabolites, including increased nucleosides and beta-oxidation intermediates. If *C. albicans* were capable of this shift in secreted metabolites within tumors, this may provide tumor cells with supplemental building blocks to recover after radiation-induced DNA damage. Future experiments will assess whether *C. albicans* behaves similarly in coculture conditions with cancer cells, as well as within murine and human tumors, to further uncover the potential feedback loop of *C. albicans* colonization and intratumoral hypoxia.

This study begins to establish a causal relationship between the presence of *Candida albicans* within rectal tumors and the development of a treatment-resistant phenotype. Given that rectal cancer is treated with chemotherapy, radiation, and surgery, gaining a stronger understanding of which patients may respond to certain therapy types would be helpful in treatment planning. Further, there may be a window to deplete intratumoral fungi, specifically *C. albicans*, with hopes of improving therapy response. Continued experimentation will assess whether the phenotype we observed is cancer-type or treatment-type specific, as we are currently building several other mouse models of cancer to test this microbe and its effects. Additionally, we plan to expand upon the rectal cancer model we currently are using to assess whether fungal dose, radiation dose, or treatment duration impacts the phenotype observed. Ultimately, understanding the complex role of specific microbes functioning within their own intratumoral niche is critical to improve outcomes in cancer patients whom treatments continue to fail.

## Supplementary Information

**Supplementary Table S1.**
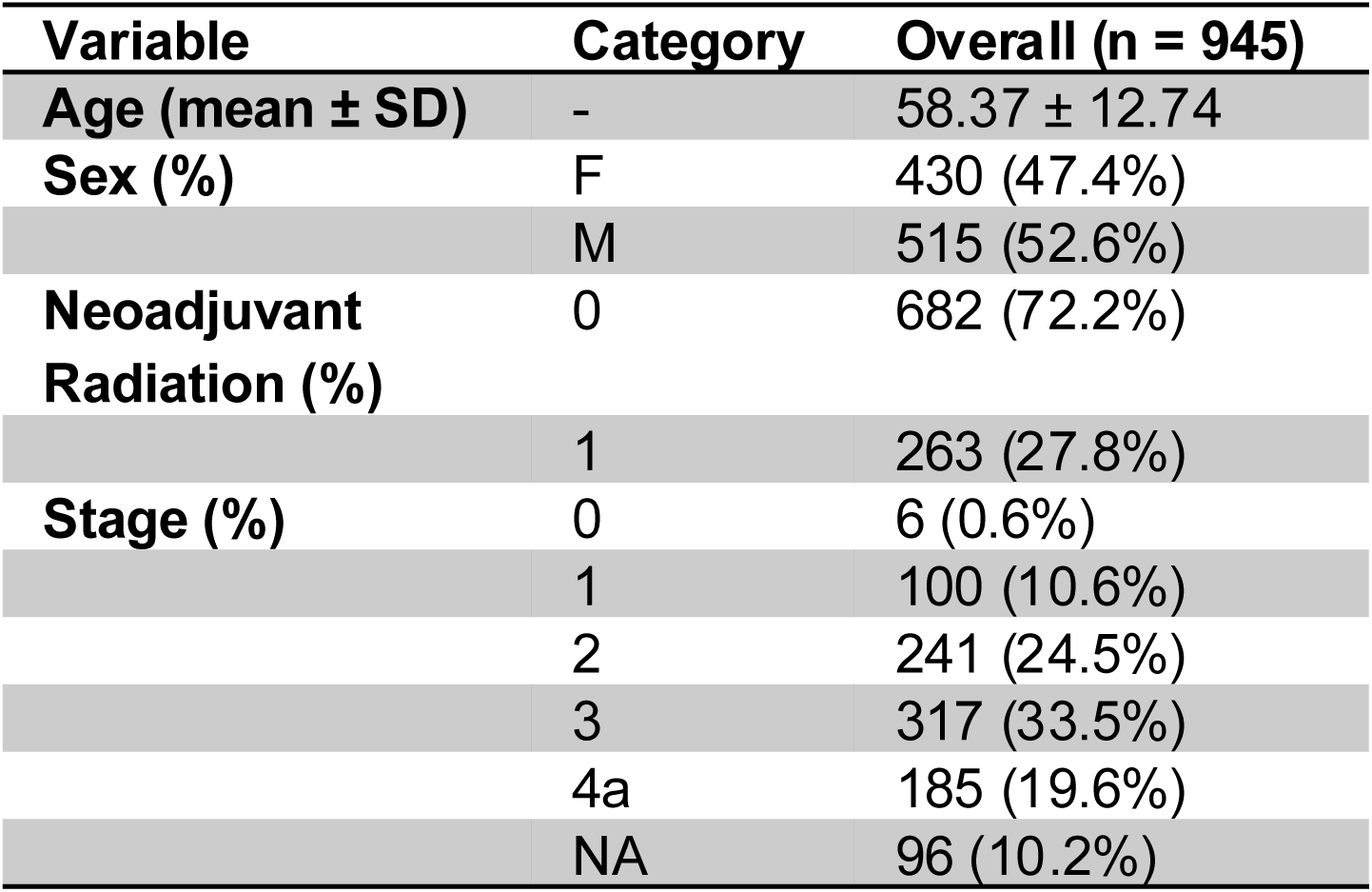
Demographics of the ORIEN cohort.

**Supplementary Table S2.** Demographic and clinical characteristics of OSU cohort.

| Variable |  | Overall (n = 30) |
| --- | --- | --- |
| Age at Enrollment (mean ± SD) | - | 61.90 ± 12.44 |
| Gender (%) | F | 14 (46.7%) |
|  | M | 16 (53.3%) |
| Stage (%) | I | 1 (3.3%) |
|  | II | 24 (80.0%) |
|  | III | 1 (3.3%) |
|  | IV | 4 (13.3%) |
| Location of Tumor (%) | Rectum | 6 (20.7%) |
|  | Sigmoid | 6 (20.7%) |
|  | Cecum | 2 (6.9%) |
|  | Ascending Colon | 1 (3.4%) |
|  | Transverse Colon | 1 (3.4%) |
|  | Descending Colon | 1 (3.4%) |
|  | Hepatic Flexure | 1 (3.4%) |
|  | Splenic Flexure | 1 (3.4%) |
|  | Rectosigmoid Junction | 1 (3.4%) |
|  | Other | 9 (31.0%) |
| Type of Treatment (%) | Surgery only | 1 (3.3%) |
|  | Radiation only | 1 (3.3%) |
|  | Chemotherapy | 22 (73.3%) |
|  | Combined therapy | 3 (10.0%) |
|  | Palliative care | 2 (6.7%) |
|  | Unknown | 1 (3.3%) |
| Current Status (%) | Alive, no disease | 20 (66.7%) |
|  | Alive, with disease | 2 (6.7%) |
|  | Deceased | 8 (26.6%) |
| AJCC Score (%) | 0 | 5 (16.7%) |
|  | 1 | 1 (3.3%) |
|  | 2 | 8 (26.7%) |
|  | 3 | 1 (3.3%) |
|  | 4 | 7 (23.3%) |
|  | 5 | 2 (6.7%) |
|  | 6 | 6 (20.0%) |

**Supplementary Table S3.**
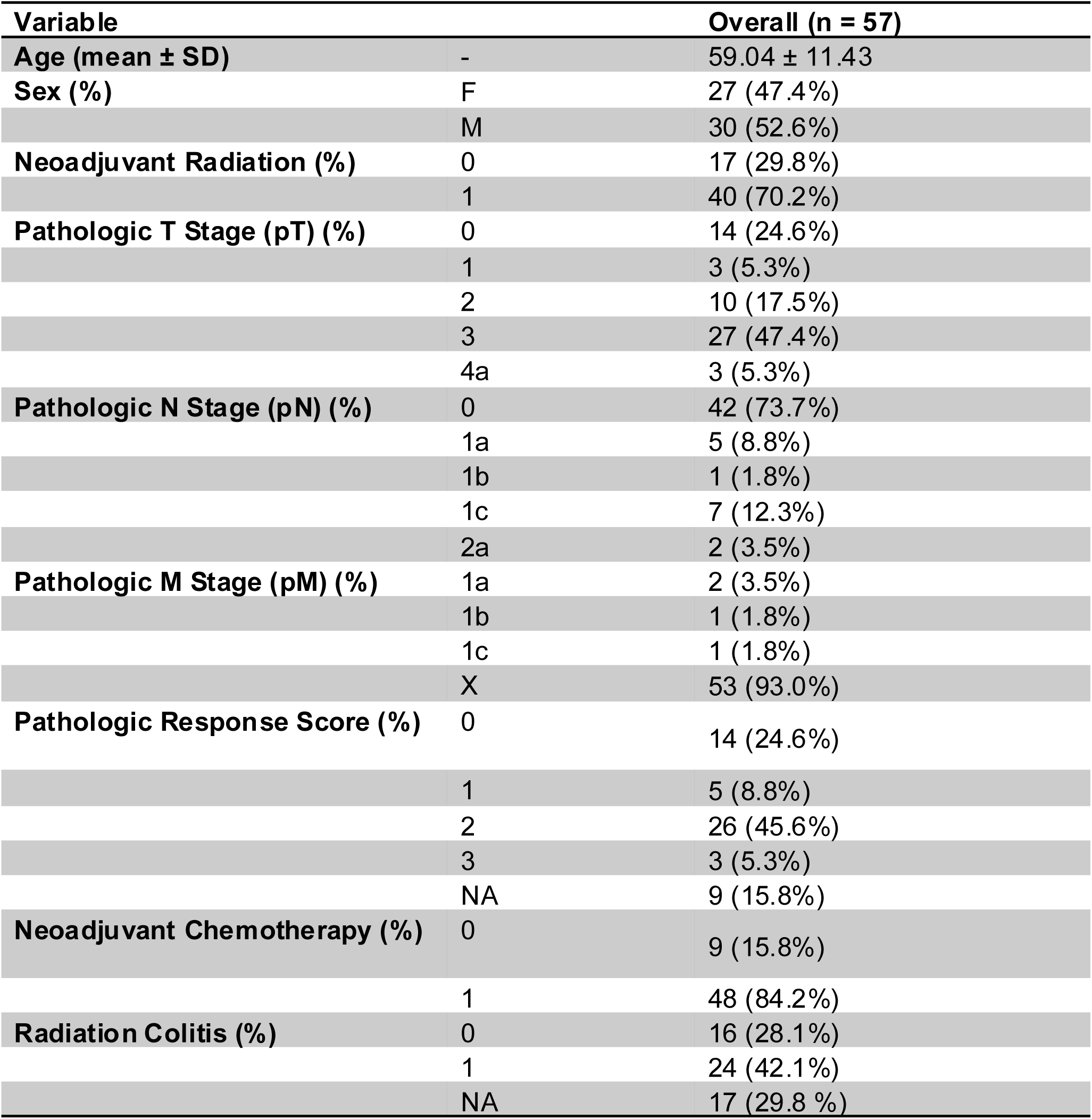
Demographic and clinical characteristics of Iowa cohort.

| Variable |  | Overall (n = 57) |
| --- | --- | --- |
| Age (mean ± SD) | - | 59.04 ± 11.43 |
| Sex (%) | F | 27 (47.4%) |
|  | M | 30 (52.6%) |
| Neoadjuvant Radiation (%) | 0 | 17 (29.8%) |
|  | 1 | 40 (70.2%) |
| Pathologic T Stage (pT) (%) | 0 | 14 (24.6%) |
|  | 1 | 3 (5.3%) |
|  | 2 | 10 (17.5%) |
|  | 3 | 27 (47.4%) |
|  | 4a | 3 (5.3%) |
| Pathologic N Stage (pN) (%) | 0 | 42 (73.7%) |
|  | 1a | 5 (8.8%) |
|  | 1b | 1 (1.8%) |
|  | 1c | 7 (12.3%) |
|  | 2a | 2 (3.5%) |
| Pathologic M Stage (pM) (%) | 1a | 2 (3.5%) |
|  | 1b | 1 (1.8%) |
|  | 1c | 1 (1.8%) |
|  | X | 53 (93.0%) |
| Pathologic Response Score (%) | 0 | 14 (24.6%) |
|  | 1 | 5 (8.8%) |
|  | 2 | 26 (45.6%) |
|  | 3 | 3 (5.3%) |
|  | NA | 9 (15.8%) |
| Neoadjuvant Chemotherapy (%) | 0 | 9 (15.8%) |
|  | 1 | 48 (84.2%) |
| Radiation Colitis (%) | 0 | 16 (28.1%) |
|  | 1 | 24 (42.1%) |
|  | NA | 17 (29.8%) |

